# Investigating functional brain network integrity using a traditional and novel diagnostic system for neurodevelopmental disorders

**DOI:** 10.1101/396317

**Authors:** Dina R. Dajani, Catherine A. Burrows, Paola Odriozola, Adriana Baez, Mary Beth Nebel, Stewart H. Mostofsky, Lucina Q. Uddin

**Affiliations:** Department of Psychology, University of Miami, Coral Gables, FL; Institute on Community Integration, University of Minnesota, Minneapolis, MN; Department of Psychology, Yale University, New Haven, CT; Center for Neurodevelopmental and Imaging Research, Kennedy Krieger Institute; Department of Neurology, Johns Hopkins University School of Medicine; Department of Psychiatry and Behavioral Sciences, Johns Hopkins University School of Medicine; Neuroscience Program, University of Miami Miller School of Medicine, Miami, FL

**Author notes:** Corresponding author: Dina R. Dajani, Department of Psychology, University of Miami, P.O. Box 248185-0751, Coral Gables, FL 33124.

**Keywords:** research domain criteria, functional connectivity, nosology, DSM, ASD, ADHD

## Abstract

**Background:** Current diagnostic systems for neurodevelopmental disorders do not have clear links to underlying neurobiology, limiting their utility in identifying targeted treatments for individuals. Several factors contribute to this issue, including the use of small samples in neuroimaging research and heterogeneity within diagnostic categories. Here, we aimed to investigate differences in functional brain network integrity between traditional diagnostic categories (autism spectrum disorder [ASD], attention-deficit/hyperactivity disorder [ADHD], typically developing [TD]) and carefully consider the impact of comorbid ASD and ADHD on functional brain network integrity in a large sample. We also assess the neurobiological validity of a novel, potential alternative nosology based on behavioral measures of executive function.

**Method:** Five-minute resting-state fMRI data were obtained from 168 children (128 boys, 40 girls) with ASD, ADHD, comorbid ASD and ADHD, and TD children. Independent component analysis and dual regression were used to compute within- and between-network functional connectivity metrics at the individual level.

**Results:** No significant group differences in within- nor between-network functional connectivity were observed between traditional diagnostic categories (ASD, ADHD, TD) even when stratified by comorbidity (ASD+ADHD, ASD, ADHD, TD). Similarly, subgroups classified by executive functioning levels showed no group differences.

**Conclusions:** Using clinical diagnosis and behavioral measures of executive function, no group differences were observed among the categories examined. Therefore, we suggest that brain imaging metrics may more effectively define clinical subgroups than behavioral metrics, and may contribute to the establishment of a neurobiologically valid nosology for neurodevelopmental disorders.

## Background

Autism spectrum disorder (ASD) and attention-deficit/hyperactivity disorder (ADHD) are behaviorally heterogeneous, prevalent neurodevelopmental disorders with few clear links between diagnostic criteria and specific neurobiological alterations^1,2^. These disorders exhibit shared deficits in executive function^3^ and associated functional brain alterations^4,5^, which may be exacerbated by the high rates of comorbidity between ASD and ADHD^6^. These challenges limit the utility of the diagnostic and statistical manual of mental disorders’ (DSM-5^7^) criteria as predictors of etiology or treatment response^8^. The NIH has proposed the Research Domain Criteria (RDoC) framework, which instead suggests the use of neurocognitive constructs that are linked to underlying neurobiology to investigate mental health disorders^9^. Following these guidelines, a potential alternative nosology may be developed that is specifically tied to targeted treatments. In this study, we first assess the neurobiological validity of traditional diagnostic categories by evaluating differences in functional brain network integrity between children with ASD, ADHD and typically developing (TD) children. Comorbidity between ASD and ADHD is rarely considered in neuroimaging studies. Here, we explicitly examine brain network connectivity in children with comorbid ASD and ADHD separately from children with ASD (without ADHD), ADHD (without ASD), and TD children. Finally, we assess the neurobiological validity of a novel classification system based on behavioral measures of executive function as an alternative to the DSM-5 classification system.

The DSM-5 defines ASD according to behavioral symptoms of social/communication deficits and restricted and repetitive behaviors; ADHD is defined by either primarily inattentive or hyperactive symptoms, or a combination of both^7^. Although there exists a large body of work implicating functional brain network alterations in ASD and ADHD^1^, to date, no biomarkers have been identified to supplement diagnosis of these disorders. Here, we focus on functional connectivity as an important marker for dysfunction in these neurodevelopmental disorders.

Case-control studies comparing functional connectivity in ASD and ADHD to TD children have produced largely inconsistent results^11,12^, although studies of children with ADHD appear to be more consistent than studies of ASD^14^. Both hyperconnectivity^15,16^ and hypoconnectivity^17,18^ have been reported in children with ADHD and ASD. The neuroscience field is becoming increasingly aware of the lack of reproducible findings across studies, which may be the result of low sample sizes, inflated false-positive rates due to analytic choices, and heterogeneity inherent to groups of interest^19^. In response, the field is calling for a focus on replication studies using adequately powered samples^20^. Here, we aim to resolve the discrepancy in prior studies characterizing brain networks of children with ASD and ADHD by comparing functional network integrity between groups in a large sample of children.

Few neuroimaging studies have directly compared network functional connectivity of children with ASD and ADHD, resulting in inconclusive findings of both common and distinct network alterations across case-control studies^4,21^. Further, only two prior functional neuroimaging studies considered the impact of comorbidity on diagnostic group differences by examining an ASD with comorbid ADHD group distinct from non-comorbid ASD and ADHD groups^4,5^. These studies demonstrated that children with ASD and comorbid ADHD exhibit functional brain network abnormalities that resemble alterations in children with ASD (without ADHD) and ADHD (without ASD) plus unique abnormalities specific to comorbidity between ASD and ADHD^4,5^. These prior results suggest that some brain network findings may be nonspecific across ASD and ADHD, and some may only hold for subgroups within a disorder (e.g., a comorbid group). The current study addresses both of these concerns in the context of two highly prevalent neurodevelopmental disorders. The first aim of this study is to test differences in functional connectivity between children with ASD, ADHD, comorbid ASD and ADHD, and TD children.

Inconsistent findings and non-specificity of functional network alterations in ASD and ADHD may also be attributed to the heterogeneity characteristic of these disorders. Individual differences among children within a DSM diagnostic category are increasingly recognized across the biological psychiatry field, with recent calls to account for this variability in research studies^1,22^. One possible approach for parsing heterogeneity in these disorders is to define more homogeneous subgroups of children with ASD and ADHD. Based on DSM-5 criteria, there are no currently defined subgroups for ASD^7^. Although ADHD has three DSM-defined subgroups (Inattentive, Hyperactive/Impulsive, and Combined), these are currently inadequate to capture the full range of symptoms^23^ or to predict treatment response^2^. In addition to high levels of within-group variability, there is considerable overlap in both phenotypic and biological alterations in ASD and ADHD, one of the most striking similarities being common difficulties in executive function^22^. Shared alterations in structural^10^ and functional^4^ neural underpinnings of executive function across ASD and ADHD have likewise been reported. Importantly, Chantiluke et al. (2014) showed that children with “pure” ASD and ADHD exhibited little to no disorder-specific functional alterations during a temporal discounting task, whereas the comorbid group exhibited pronounced functional differences compared with all other groups^5^. This finding highlights the inadequacy of defining subgroups solely within a single disorder, and instead calls for the definition of subgroups that may cut across disorders.

To this end, we recently leveraged both theoretical (focusing on executive functions) and data-driven (using a subgrouping method called latent profile analysis) computational psychiatry approaches to develop a possible alternative categorization of neurodevelopmental disorders. We demonstrated that behavioral measures of executive function can be used to define subgroups of children across various diagnostic groups: ASD, ADHD, comorbid ASD and ADHD, and TD children^24^. Three subgroups emerged— “above average,” “average,” and “impaired” executive functions— which crossed traditional diagnostic boundaries. Here, we follow up these findings by assessing the neurobiological validity of the current DSM categories of ASD and ADHD, in addition to considering comorbidity. Moreover, we assess the neurobiological validity of an alternative nosology based on executive function subgroups (i.e., “above average,” “average,” and “impaired” subgroups). We operationalize neurobiological validity as the separability of groups based on within- and between-network functional connectivity of three major intrinsic connectivity networks (ICNs) important for cognition^27^: the frontoparietal network (FPN), salience network (SN), and DMN^28^.

We predicted that children with ADHD would exhibit reduced DMN connectivity and stronger DMN-FPN and DMN-SN coupling^14^ compared with TD children. We expected that children with ASD would exhibit hyperconnectivity within FPN, DMN, and SN^15,29^ compared with TD children. Inconsistent findings comparing ASD, ADHD, and ASD with comorbid ADHD groups precludes establishing well-founded *a priori* hypotheses specific to the comorbid ASD and ADHD group. Further, we anticipated that there would be a parametric increase in functional connectivity of the FPN and SN across executive function subgroups, such that the “impaired” subgroup would exhibit the lowest functional connectivity and the “above average” group would exhibit the highest functional connectivity^30^.

## Methods and Materials

### Participants

Participants aged 8 to 13 years (*N=*168) included a subset of children used in our previous study^24^. Written informed consent was obtained from all legal guardians and written assent was obtained from all children. All procedures were approved by the Institutional Review Board at the Johns Hopkins School of Medicine and all methods were carried out in accordance with the approved guidelines.

### Diagnostic and IQ measures

Community diagnoses of ASD were confirmed with the Autism Diagnostic Observation Schedule (ADOS-G^32^ or ADOS-2^33^, based on study enrollment date) and Autism Diagnostic Interview-Revised (ADI-R^34^). The Diagnostic Interview for Children and Adolescents IV^35^ was used to confirm community ADHD diagnoses, determine whether children with ASD had comorbid ADHD, and for exclusionary purposes. Community diagnoses of ADHD were also confirmed with the Conners’ Parent Rating Scales (CPRS-R:L^36^ or CPRS-3^37^, based on study enrollment date) and the ADHD Rating Scale IV, Home version^38^ (Table 1). Executive functions used for the latent profile analysis were measured primarily with a parent-report (eight subscales of the BRIEF^40^), in addition to two laboratory measures, the Statue subscale of the NEPSY-II^41^ and the backward digit span of the WISC-IV. See supplementary information for more details.

**Table 1.**
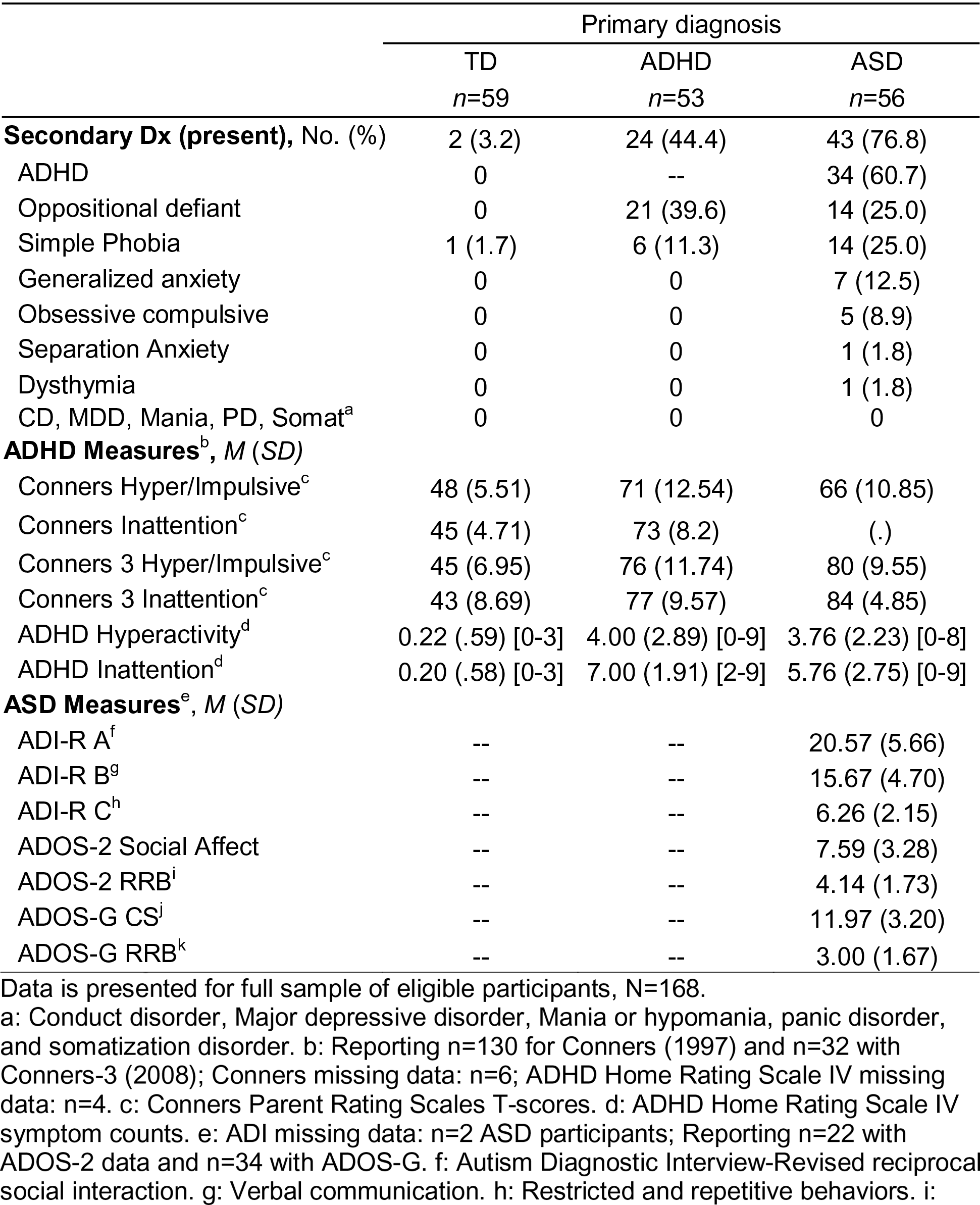
Diagnostic information.

### Subsamples

To compare the integrity of ICNs between groups delineated by primary diagnosis (ASD, ADHD, TD), three equally sized diagnostic groups (*N*=129, three groups of *n*=43) were randomly selected from the larger sample of 168 participants in order to produce unbiased group ICA results (hereafter, ‘Diagnostic group sample’). These diagnostic groups were delineated by *primary* diagnosis; individuals may have had additional *secondary* diagnoses. For example, the “ASD” group included individuals with comorbid psychiatric diagnoses, including ADHD (*n*=26, 61%). Some children in the “ADHD” group had comorbid psychiatric diagnoses, but this *did not* include comorbid ASD, given that ASD is commonly considered a primary and not secondary diagnosis (see Table 1 for more details). Diagnostic groups did not differ in age, sex, handedness, FSIQ, or head motion (mean framewise displacement [FD^42^], translational and rotational motion, Table 2).

**Table 2.**
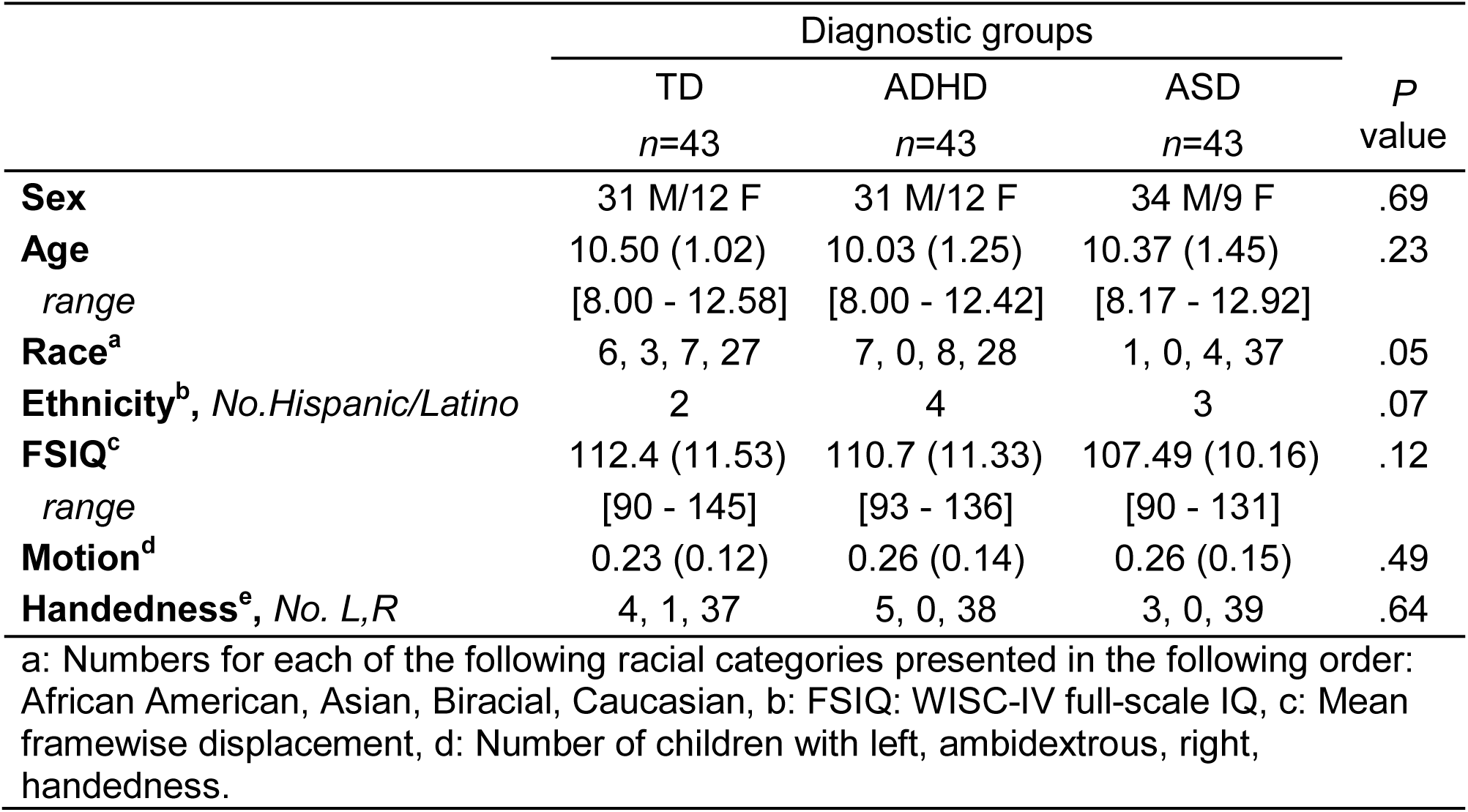
‘Diagnostic group sample’ demographics

To address the issue of comorbidity in assessing diagnostic group differences, an additional group analysis was performed. Diagnostic groups were categorized as TD, ADHD, ASD (without ADHD), and ASD+ADHD (ASD with comorbid ADHD). Due to sample size limitations, group size for this analysis was limited to 22 (*N*=88, ‘Diagnostic comorbid sample’). These groups did not differ in age, sex, handedness, or head motion (Table S1). Due to sample size limitations, we were unable to match the diagnostic comorbid samples on FSIQ (*p*<.001).

A subset of the 168 participants eligible for this study was generated to ensure equal group sizes for each executive function (EF) subgroup (‘EF subgroup sample’, *N*=129, three groups of *n*=43). The ‘EF subgroup sample’ was representative of the *Dajani* et al. (2016) sample^24^ in EF scores, age, FSIQ and the distribution of diagnostic categories within each EF subgroup. EF subgroups did not differ in age, sex, handedness or mean head motion, but did differ on FSIQ (*p*<.001) and reached a near-significant difference in mean FD for the raw data (*p*=.06, Table S2). Following preprocessing, there were no group differences in mean FD (*p*=.36, Figure 2). The difference in FSIQ was expected given the subgroups were delineated based on EF, which tends to be highly correlated with IQ metrics^43^.

**Figure 1.**
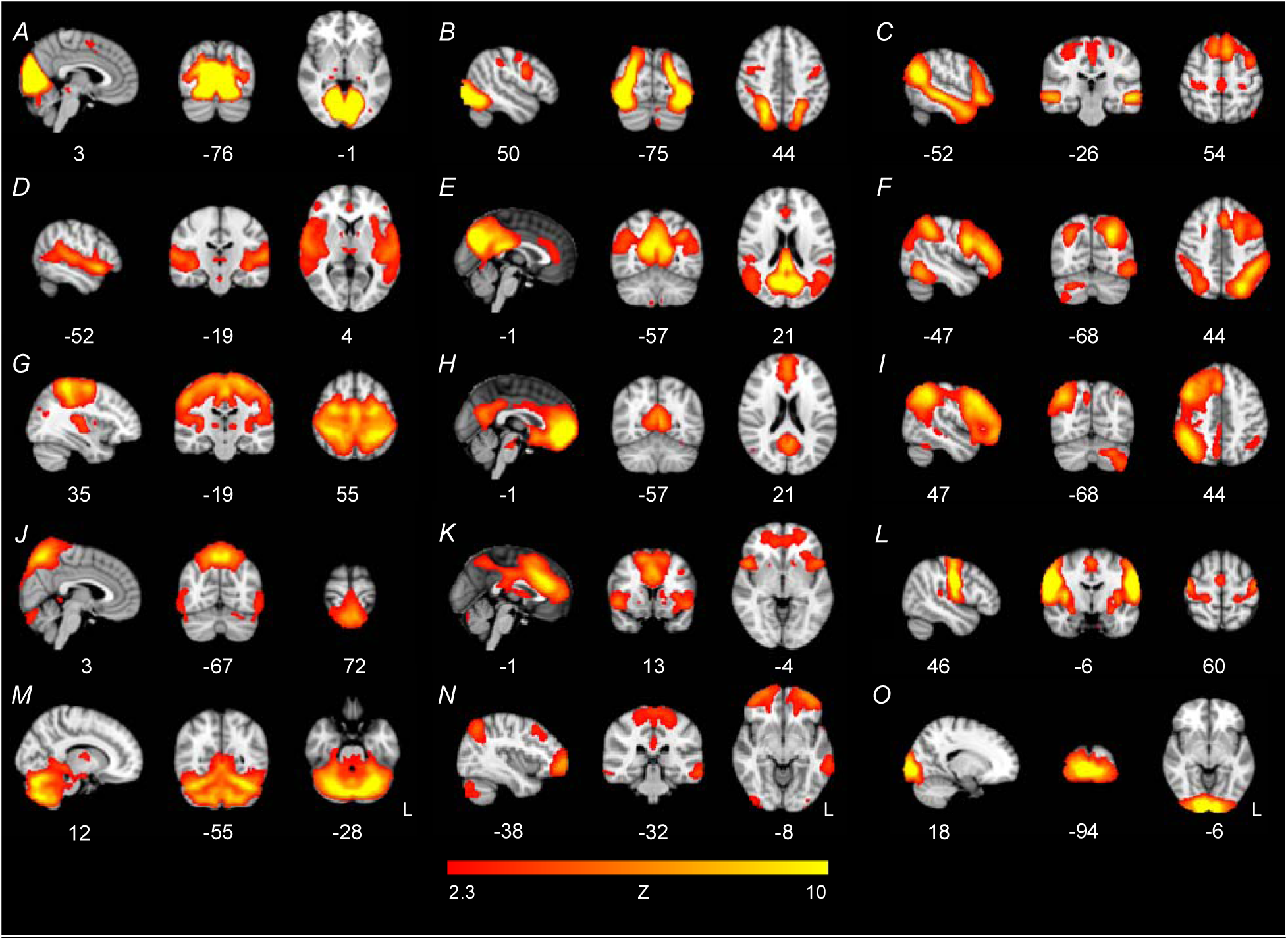
Fifteen-component ICA results for the ‘EF subgroup’ sample. Components E (posterior DMN), F (L FPN), H (anterior DMN), I (R FPN), and K (SN) were used to assess group differences in network connectivity. One artifactual component emerged (component J).

**Figure 2.**
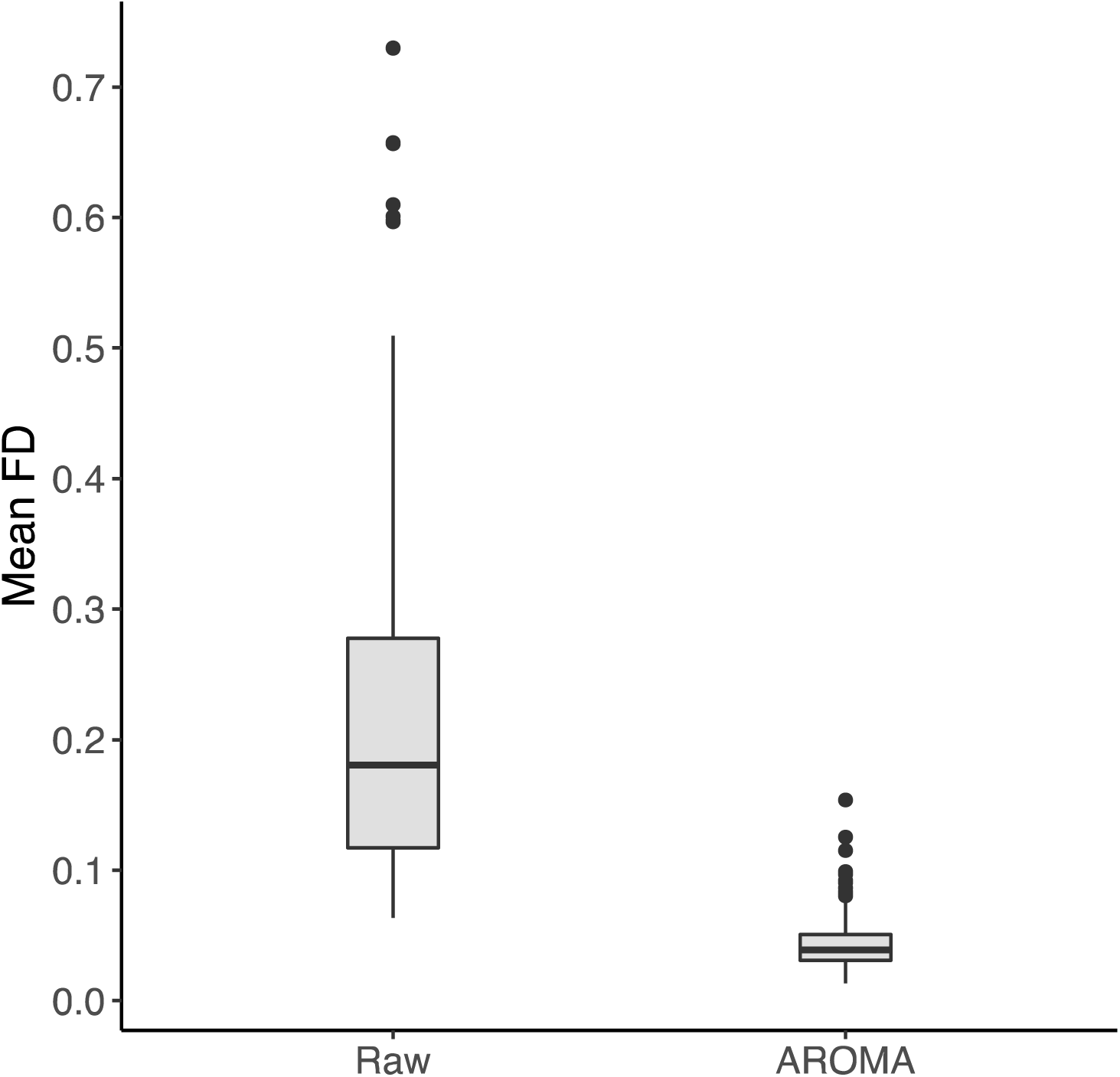
Effect of rigorous motion correction. Mean FD is in mm, displayed on raw data and data after preprocessing with ICA-AROMA

### Data acquisition

Resting state fMRI (rs-fMRI) data were acquired for participants on a Phillips 3T scanner using an 8-channel head coil (TR=2.5s, flip angle=70°, sensitive encoding acceleration factor=2, 3mm slices, voxel size= 2.7×2.7×3 mm). Most participants had a 156-volume dataset, but a subset had a shorter acquisition time of 128 volumes (156 volumes: *n*=114, 128 volumes: *n*=15 in the ‘EF subgroup sample’). High-resolution T1-weighted scans were also acquired to facilitate registration of the functional image to standard space (TR=8.0ms, TE=3.7ms, 1mm isotropic voxels). Participants were asked to withhold stimulant medication (e.g., Adderall) the day before and on the day of MRI scanning, similar to prior neuroimaging studies comparing children with ASD and ADHD^4,44^. Non-stimulant medications were continued as prescribed (e.g., antidepressants, allergy medication, see Table S3 for detailed medication status information).

### Experimental Design and Statistical Analysis

#### Preprocessing

Dual regression analyses employed in this study require all participants to have equal rs-fMRI scan lengths, therefore all participants’ rs-fMRI data were truncated to the shortest participant’s scan length by removing volumes at the end of the scan, resulting in all scans including 121 volumes (5 minutes of data). Standard preprocessing was conducted in FSL 5.0.9. In addition, a state-of-the-art ICA-based denoising procedure (ICA-AROMA^45^) was used to remove motion-related artifacts in native space (see supplemental information).

#### Group ICA

We ran a group ICA using the FSL MELODIC v3.14 toolbox (https://fsl.fmrib.ox.ac.uk/fsl/fslwiki/MELODIC) with temporal concatenation to identify common spatial patterns across participants. Five components of interest were manually identified from the group ICA by two of the authors (DRD and LQU): right FPN^50^, left FPN^50^, SN^51^, and anterior and posterior DMN^50^. To ensure unbiased networks were produced for each subsample and subsequent analyses, a separate group ICA model was run for each of the three subsamples. Networks were qualitatively highly similar across subsamples (Figure S1).

#### Dual Regression

Dual regression is a reliable technique which allows for the identification of group differences in the spatial and temporal features of ICNs common to the entire sample^25,26^. Using FSL’s dual regression command (https://fsl.fmrib.ox.ac.uk/fsl/fslwiki/DualRegression/UserGuide), individual-level spatial and temporal maps were constructed for each of the five components of interest generated from the group ICA. To test group differences in within-network connectivity, the normalized individual-level spatial maps were subjected to permutation testing using FSL’s randomise tool for each of the five components of interest (https://fsl.fmrib.ox.ac.uk/fsl/fslwiki/Randomise/UserGuide, 5000 permutations). To test for group differences in between-network connectivity, the FSLNets package was implemented in MATLAB (https://fsl.fmrib.ox.ac.uk/fsl/fslwiki/FSLNets). For each subsample, *F*-tests were conducted to ascertain the presence of group differences in within-network and between-network connectivity strength. Significance was determined using a threshold-free cluster enhancement of *p*<.05 (FWE-corrected) for both within- and between-network analyses, as in previous studies^15,29^.

## Results

### Effect of rigorous motion correction

A 2×1 repeated-measures ANOVA was used to assess whether motion (indexed by mean FD^42^) decreased as a function of preprocessing with ICA-AROMA by comparing mean FD for raw and preprocessed data. The ANOVA demonstrated a significant decrease in mean FD following preprocessing, *F*(1, 332)=25.36, *p*<.001 (raw: *M*=.22 *SD*=.14, preprocessed: *M*=.04 *SD*=.02, Figure 2).

### Comparisons between diagnostic groups

No within- or between-network connectivity differences were found when comparing diagnostic groups (TD, ADHD, ASD; within-network: FWE-corrected *p*’s>.26, between-network: FWE-corrected *p*’s>.37). When comparing diagnostic groups and statistically controlling for the effects of EF subgroup, *F*-tests remained non-significant (within-network: FWE-corrected *p*’s>.20, between-network: FWE-corrected *p*’s>.11). *Comparisons between comorbid diagnostic groups*

When diagnostic groups were broken down according to comorbidity (TD, ADHD [without ASD], ASD [without ADHD], ASD+ADHD), *F*-tests revealed no group differences in within-network (FWE-corrected *p*’s>.30) or between-network connectivity (FWE-corrected *p*’s>.14).

### Comparisons between EF subgroups

Group differences in network connectivity strength were assessed across five ICNs between children with “above average,” “average,” or “impaired” EF. *F*-tests revealed no significant differences between EF subgroups in within-network nor between-network connectivity (within-network: FWE-corrected *p*’s>.24, between-network: FWE-corrected *p*’s>.29). Likewise, after controlling for diagnostic status, no differences in connectivity emerged (within-network: FWE-corrected *p*’s>.25, between-network: FWE-corrected *p*’s>.47).

## Discussion

With the recent exponential increase in computational power and growing awareness of the limitations of current psychiatric diagnostic systems, there have been numerous recent calls to leverage the strengths of computational psychiatry approaches to develop a more parsimonious and neurobiologically valid nosology of mental health disorders^52,53^. This study is the first to use ICA dual regression, a reliable data-driven approach to investigate differences in both within- and between-network functional connectivity between clinical groups, to assess the neurobiological separability of a traditional diagnostic classification system and a novel subgrouping system based on behavioral measures of executive function, while rigorously correcting for motion-related artifacts that are pervasive in pediatric psychiatric populations. We also carefully consider the impact of comorbidity between ASD and ADHD on brain network functional connectivity. Results indicate that the current DSM categories classify children into groups exhibiting negligible functional connectivity differences of major cognitive networks, suggesting limited neurobiological validity of current DSM categories. Contrary to our hypotheses, executive function subgroups displayed limited differences in functional connectivity of major cognitive networks, suggesting that a nosology based solely on a behavioral subgrouping system may not map onto differences in underlying neurobiology.

Comparisons between traditional diagnostic categories (i.e., ASD, ADHD, and TD), which are based on observable symptoms according to DSM criteria, showed no differences in functional connectivity across major cognitive networks. Several reasons may explain these null findings. First, it may be that these diagnostic categories do exhibit differences in functional connectivity, but findings may be method-specific and do not extend to dual regression ICA-based analyses. Second, it is possible that a moderating factor, such as sample-specific IQ, gender distribution or symptom severity, may affect whether differences in functional connectivity are observed. Third, previously published findings may simply represent false positives. Finally, it may be that this study represents a false negative.

Prior studies comparing TD children and children with ASD and ADHD using ICA-derived networks report network connectivity differences between diagnostic groups. Specifically, studies of individuals with ADHD report reduced segregation of DMN-FPN networks in children, adolescents, and adults^54,55^, but results from within-network connectivity studies are less coherent. For example, both hyperconnectivity^16^ and hypoconnectivity^18^ have been reported in ADHD for the FPN and the DMN. The ASD literature is similarly inconsistent, even when limited to studies using dual regression ICA. For example, there have been reports of both within-network hyperconnectivity^15,29^ and hypoconnectivity of the FPN^17^ and DMN^17,56^. Although results are not consistent across studies, they clearly demonstrate that positive results are possible when using ICA dual-regression to investigate network integrity in children with ASD and ADHD, ruling out the possibility that previously reported network connectivity differences in ASD and ADHD are method-specific. Discrepancies in results across studies may be due to individual differences in network connectivity within these traditional diagnostic categories, thus diminishing their neurobiological separability, and ultimately, their validity.

Heterogeneity within ASD and ADHD categories, characterized by a wide range of IQ, symptom severity, and the presence or absence of comorbid psychiatric disorders, may lead to inconsistent findings across studies, ultimately culminating in non-generalizable results. Our null findings when comparing diagnostic groups are in line with a recent review of classification studies of ASD and ADHD using neuroimaging techniques, which suggested that heterogeneity within and between disorders may be limiting researchers’ (and machines’) ability to distinguish disorders based on neurobiology^57^. Two factors may be contributing to the limited neurobiological separability of ASD and ADHD categories: high within-category heterogeneity and high between-category similarity. There is considerable evidence for the existence of subgroups within ASD^58^ and ADHD diagnostic categories^59^, which are often delineated with neuropsychological measures. Recent evidence has also emerged for neurobiological subgroups of ADHD based on network connectivity of the reward system^60^ and other large-scale brain networks^61^. In addition, there is substantial overlap in ASD and ADHD categories in symptomatology (e.g., social skills deficits), behavioral domains beyond symptoms (e.g., executive dysfunction), and genetic factors^62^ in addition to high rates of comorbidity between ASD and ADHD. In sum, these diagnostic categories are not distinct from one another based on numerous criteria. Evidence for both high within-group heterogeneity and between-group overlap reduces the validity of current DSM categories for ASD and ADHD, and thus may explain our lack of dissociability between disorders based on functional connectivity metrics.

Nonetheless, the results presented here contradict past findings of significant differences in functional connectivity between children with ASD, ADHD and TD children, which may be explained by the improved methodology used in this study. Past studies using ICA dual regression to compare children with ASD or ADHD to TD children used small sample sizes (n=20-25 per group)^15,18,29,56,63,64^. It is well established that low sample sizes contribute to reduced power to detect true results. However, it is less appreciated that positive results reported from underpowered studies *also* have a greater likelihood of being false^19,65^. Further, the exorbitant number of researcher degrees of freedom available in fMRI analyses, including choice of preprocessing pipeline, analysis method, group matching procedures, thresholding procedures, and multiple comparison correction lead to inflated false-positive rates^19^. Here, we nearly double the sample size of past studies and arrive at null results, which in combination with the fact that past results are inconsistent across studies, suggests that some previous results could be attributed to false positives^65^. In addition, we employ a rigorous motion correction procedure not used in any previous study of functional connectivity differences in ASD and ADHD. ICA-AROMA is a denoising approach that reduces the likelihood of group differences emerging solely due to differences in in-scanner motion. The strength of long-range functional connectivity is underestimated in cases of increased head motion^66^, which is rife in studies of youth with neurodevelopmental disorders^46^. Previous studies of within-network connectivity of the DMN in both children with ASD^56,64^ and ADHD^14^ that did not use ICA-AROMA may have been influenced by residual effects of motion. Taken together, these results suggest that findings of reduced long-range connectivity in ASD and ADHD may be a byproduct of increased motion artifacts in these clinical groups.

Few neuroimaging studies consider the impact of comorbid ASD and ADHD on case-control findings, and instead rely on the primary diagnosis of children to classify patients. This leads to inherent heterogeneity in clinical groups studied, likely leading to inconsistencies in findings across the neuroimaging literature. The current study is one of the first functional neuroimaging studies to directly compare children with ASD, ADHD, comorbid ASD and ADHD, and TD children, and the first to do so using ICA dual regression combined with ICA-AROMA. Here, we do not find statistically significant differences between ASD and ADHD groups, even when accounting for comorbidity.

Compared to the group difference tests for the traditional diagnostic groups (ASD, ADHD, TD) and executive function subgroups, the analysis accounting for comorbidity may have been underpowered, with only 22 participants per group. Therefore, it is possible that the lack of observed differences in functional connectivity in children with comorbid ASD and ADHD represent a false negative. Ideally, this analysis should be replicated in future studies with a larger sample size to confirm whether group differences in functional connectivity exist between children with comorbid ASD and ADHD and other clinical and typically developing populations.

As an alternative nosology to DSM-5 classifications, we tested whether there existed differences in functional connectivity between “above average,” “average” and “impaired” EF subgroups^24^. Contrary to our hypotheses, we observed no differences in functional connectivity between these subgroups, suggesting that EF-defined subgroups are not neurobiologically distinct based on networks important for cognition. There are multiple possible explanations for this null result. One possibility is that EF in children may be best assessed dimensionally rather than categorically^67^. The primary goal of this study was to assess the validity of the traditional categorical nosology (i.e., the DSM-5) and an alternative categorical nosology (i.e., EF subgroups). In characterizing neurodevelopmental and psychiatric disorders, recent studies have revealed the importance of testing whether categorical, dimensional, and hybrid categorical-dimensional models best describe psychiatric disorders^67^. Future studies would benefit from testing both symptom-based and executive function-based dimensional models to determine whether these may better characterize neurobiological differences between children. Additionally, our results may have been influenced by our choice of measures for EF subgrouping, which were primarily based on a parent-report of EF symptoms (i.e., the BRIEF^40^). We chose to focus on a parent-report of EF symptoms because of the strong psychometric properties of the measure and ease of obtaining the information in a clinical setting. Although using a parent-report of observable EF clearly has easily translatable implications for clinical practice, which is a current challenge for the field^68^, the choice to focus on a parent-report of behavior to characterize psychiatric disorders may be many steps removed from the underlying neurobiology. This leads to a rather simple explanation for these results – that behavior does not map one-to-one to underlying neurobiology.

There are numerous lines of evidence to suggest that brain-behavior relationships are not simply one-to-one^69^. Varying types of functional network miswirings across development may manifest as a singular phenotype^70^, suggesting that distinct brain abnormalities may appear behaviorally as the same neurodevelopmental disorder. Likewise, disparate genetic etiologies may lead to similar behavioral profiles^10,71^. Diagnostic categories should necessarily define neurobiologically homogeneous groups to allow for the development of targeted treatments specific to a neurobiological signature of the disorder. This suggests that advances in mental health research necessarily rely on characterizing underlying neurobiological mechanisms of pathophysiology, and that psychiatry may be fundamentally limited as long as assessments are limited to observable phenomena^72^.

Applying principles of computational psychiatry to clinical research has the potential to transform the mental health field from the current trial-and-error choice of treatments towards precision medicine. Current psychiatric diagnostic systems rely on observable behaviors to classify disease, with unknown links to underlying neurobiology^52^. Biomarkers, on the other hand, may provide information that may be able to stratify current diagnostic categories or replace symptom-based classification systems completely. Here, we aimed to leverage the strengths of computational psychiatry methodology to propose an alternative nosology based on behavioral measures of executive function, which we predicted would lead to neurobiologically distinct subgroups. Contrary to predictions, we found that EF subgroups could not be distinguished based on within- or between-network connectivity metrics of major cognitive networks, emphasizing the notion that nosologies reliant on behavioral data alone may not lead to discovery of neurobiologically distinct categories, limiting their utility in predicting prognosis and efficacious treatments.

### Limitations

Although the present study had numerous strengths including a larger sample than previous similar studies, the results reported here should be considered in light of several limitations. It is difficult to draw strong conclusions from null results, as it is possible that they may be due to under-powered samples. Children in the clinical samples had various psychiatric comorbidities aside from ASD and ADHD (most commonly, oppositional defiant disorder [ODD]). While this is expected for children with ADHD given the high rates of comorbidity with ODD^73^, it may have introduced additional confounds that were not taken into account in this study. Future studies may consider the impact of different types and number of comorbid disorders on brain network integrity. One alternative explanation for our finding of no group differences in functional network integrity between EF subgroups is that another RDoC domain, such as social communication, may be better suited for developing an alternative nosology for children with neurodevelopmental disorders. We look forward to future studies exploring the efficacy of various domains and units of analysis outlined by the RDoC matrix.

### Conclusions

We present findings that traditional diagnostic categories of ASD and ADHD could not be distinguished from one another or from TD children based on within- and between-network functional connectivity of three major cognitive networks: the frontoparietal, salience, and default-mode networks. Likewise, EF subgroups did not reflect neurobiologically distinct subgroups. Our study provides evidence that behavioral variables may be less informative than initially thought to create neurobiologically informed subgroups. The field of computational psychiatry is nascent^53^, and will therefore require the pursuit of new avenues and the continuous refinement and validation of new hypotheses. Based on the current findings, we suggest future work should employ data-driven approaches applied to *neurobiological variables*, such as functional connectivity metrics, to create parsimonious, biologically validated categories of neuropsychiatric diseases.

## Acknowledgements

We thank Jeanette Mumford for her contribution to the statistical modeling of the group difference tests. This work was supported by the National Institute of Mental Health (R01MH107549) to LQU. This work was also funded by Autism Speaks and NIH: R01NS048527, R01 MH085328, R01 MH078160, the Johns Hopkins University School of Medicine Institute for Clinical and Translational Research, and NIH/NCRR CTSA Program, UL1-RR025005, to S.H.M.

## Supplementary Information

### Participants

In our previous study^1^, a latent profile analysis using eight behavioral indicators of EF was conducted to determine subgroups of children (8-13 years, *M*=10.01, *SD*=1.37, *N*=321) with varying levels of EF ability in a mixed group of typically developing children (TD, *n* = 128), children with ADHD (*n* = 93), children with ASD without ADHD (*n* = 30), and children with comorbid ASD and ADHD (*n* = 66). Three classes emerged that did not reproduce diagnostic categories: “above average”, “average”, and “impaired” EF.

*Diagnostic group sample IQ matching*. IQ was measured with the Wechsler Intelligence Scale for Children IV (WISC-IV^39^). To match diagnostic groups on FSIQ, first a subsample was generated from the sample of 168 participants by excluding participants with FSIQ < 90 and randomly excluding a proportion of TD participants with high IQs (>115). Second, participants were randomly selected to ensure each diagnostic group had equal sample sizes.

### Diagnostic measures

Community diagnoses of ASD and ADHD were confirmed with a mixture of parent report, parent interviews, and child interviews.

*ADOS-G and ADOS-2*. The Autism Diagnostic Observation Schedule-Generic (ADOS-G^2^) and the Autism Diagnostic Observation Schedule-2 (ADOS-2^3^) are child clinical interviews with structured and semi-structured parts that focus on social and communicative behaviors in children. Participants recruited prior to the release of the ADOS-2 received the ADOS-G. Community diagnoses of ASD were confirmed by a score of ≥ 7 for the total score on the ADOS-2 or the communication and social interaction score on the ADOS-G.

*ADI-R*. The Autism Diagnostic Interview-Revised (ADI-R^4^) is a parent clinical interview that assesses key diagnostic features of autism like reciprocal social interaction, communication, and repetitive/stereotyped behaviors. All ASD participants met criteria for ASD based on established cutoffs (≥ 10 for social interaction, ≥ 8 for communication/language, ≥ 3 for RRBs) except for one participant. This participant met criteria for ASD based on the social interaction and RRB subscales, but scored a 4 on the communication/language subscale. This participant was retained in the ASD group because they met criteria based on the ADOS-G (total score: 16).

*Conners’ PRS*. The Conners’ Parent Rating Scales-Revised, Long Version (CPRS-R:L^5^) and the Conners’ Parent Rating Scales-3rd Edition, Full-length (CPRS-3^6^) are parent reports of their child’s ADHD symptoms, oppositional defiant disorder and conduct disorder. T-Scores of 60 or higher on the DSM-IV (for the Revised version) and DSM-IV-TR (for the 3rd Edition) Hyperactive/Impulsive or Inattentive scales were used to confirm community ADHD diagnoses. One ADHD participant did not meet these criteria (Hyperactive/Impulsive: 58, Inattentive: 55), but did meet criteria based on the DICA-IV. TD participants who had T-Scores >65 on either the Hyperactive/Impulsive or Inattentive scales were excluded from the study.

*ADHD-RS-IV, Home version*. The ADHD Rating Scale-IV, Home version^7^ is a parent report of their child’s ADHD symptoms over the previous 6 months. Community diagnoses of ADHD were confirmed if children met for 6 out of 9 DSM-IV symptoms on either the Hyperactive/Impulsive or Inattention scales. Ten ADHD participants did not meet these criteria, but did meet criteria for ADHD based on both the DICA-IV and Conners’ PRS. TD participants who met on 4 out of 9 symptoms on either the Hyperactive/Impulsive or Inattention scales were excluded from the study.

DICA-IV. The Diagnostic Interview for Children and Adolescents IV^8^ is a structured parent interview to evaluate current psychiatric diagnoses in the child. The Attention Deficit Disorder subscale was used to: 1) confirm community ADHD diagnoses and 2) to determine whether the children with community ASD diagnoses also had comorbid ADHD. TD children who met criteria for any disorder were excluded except for one participant who met criteria for a simple phobia of bugs. Scores on the DICA-IV and clinical judgment by a child neurologist (S.H.M) determined ADHD subtype.

### Rs-fMRI data acquisition

Children completed a mock scanning session prior to fMRI data collection to acclimatize them to the scanning environment. Following data acquisition, the first 10 volumes of rs-fMRI datasets were immediately discarded to account for magnet stabilization. Children were asked to relax and focus on a crosshair while remaining as still as possible.

### Preprocessing

To be eligible for the current study, participants were required to have a good quality resting-state fMRI dataset based on visual inspection. Data quality was visually inspected for both raw structural and functional images, and following brain extraction and registration. All data included passed quality checks at each step, except for a single structural image with considerable ringing due to motion. In this case, registration passed our quality check, therefore we opted to retain this case in the final sample.

Participants with excessive in-scanner motion (> 5mm of absolute maximum motion) were excluded from the study. To reduce motion artifacts, volumes in raw rs-fMRI data that contained motion spikes (>3mm or degrees) at the beginning or end of the scan were deleted for 5 participants (volumes deleted were restricted to the first or last volumes of the scan to preserve the temporal continuity of the time series).

Standard preprocessing procedures included the following: First, structural images were brain extracted using FSL’s BET tool. Using FEAT, fMRI data underwent motion correction, 4D intensity normalization, smoothing with a 6mm kernel, and estimation of linear and non-linear warping parameters to normalize to the MNI152 2mm template. Following the removal of motion-related signals in native space using ICA-AROMA (see below for more details), warping parameters were applied to denoised functional images.

### ICA-AROMA

Accounting for head motion is especially important in rs-fMRI studies of pediatric and clinical populations^42,46^. Therefore, we employed a state-of-the-art method called ICA-AROMA^45,47^. ICA-AROMA takes advantage of ICA’s ability to separate noise from signal components at the individual level, automatically identifies head motion-related components trained on four theoretically identified features, and regresses the variance associated with these motion-related features from the individual-level data. Specifically, ICA-AROMA identifies components strongly associated with head motion estimates, high-frequency content, and brain edge and ventricle intensity. A validation study using samples of healthy controls and children with ADHD demonstrated that this method effectively removes motion-related variance from fMRI data, distance-dependent effects of motion, and preserves signals of interest^45^. In addition, ICA-AROMA has benefits over traditional motion correction procedures such as motion scrubbing^42^ or spike regression^48^ in that ICA-AROMA largely preserves the temporal degrees of freedom and autocorrelation structure in the data, and does not require model training or hand-labeling of components as required for ICA-FIX^49^. Of note, this method does not identify components related to white matter or cerebrospinal fluid, but these components were identified at the group-level ICA and removed from further analysis (see ‘Group ICA’ section).

### Group ICA

Multiple group ICAs were run specifying various component numbers (13,15, and 20) to determine the optimal number of components for this denoised dataset. ICA-AROMA removes many artifactual components prior to group ICA^9^. In the Pruim et al. (2015) paper, implementing ICA-AROMA prior to group ICA resulted in only 11 automatically estimated components compared to simply regressing motion parameters, which resulted in 22 estimated components. For this dataset, estimating fifteen components was optimal in identifying large-scale networks, while avoiding lumping of multiple networks together or dividing networks into individual brain regions (Figure 1). Of the fifteen components estimated for the ‘EF class sample’, only one component (Figure 1, component J) was identified as noise due to cerebrospinal fluid signal.

Five components of interest were manually identified from the group ICA by two of the authors (DRD and LQU): right FPN^50^, left FPN^50^, SN^51^, and anterior and posterior DMN^50^. Specifically, right and left FPN components were identified by the presence of lateralized dorsolateral PFC, ventrolateral PFC and lateral parietal cortices^51^. The SN was identified by the presence of anterior insula and dorsal ACC^51^. In children, the DMN tends to decompose into anterior and posterior components^15,29^. The anterior DMN was identified by the primary presence of ventromedial PFC and the posterior DMN was identified by the precuneus and posterior cingulate cortex^50^.

### FSLNets

FSLNets takes the individual-level time courses produced from the dual regression analysis and computes between-network correlations, resulting in a Fisher’s r to z transformed correlation matrix. Full (zero-order) correlation matrices were restricted to the 5 components of interest, resulting in a 5×5 matrix produced for each individual.

**Table S1.** ’Diagnostic comorbid sample’ demographics

|  | Diagnostic groups |  |  |  | *P* value |
| --- | --- | --- | --- | --- | --- |
|  | TD<br>*n*=22 | ADHD<br>*n*=22 | ASD<br>*n*=22 | ASD+ADHD<br>*n*=22 |  |
| **Sex** | 19 M/3 F | 14 M/8 F | 17 M/5 F | 19 M/3 F | .21 |
| **Age** |  |  |  |  |  |
| *range* | [8.00 - 12.58] | [8.08 - 12.33] | [8.00 - 12.50] | [8.00 - 12.50] |  |
| *mean* | 10.41 (1.23) | 10.17 (1.16) | 9.69 (1.30) | 10.33 (1.28) | .23 |
| **Race<sup>a</sup>** | 2, 2, 3, 15 | 6, 0, 1, 15 | 0, 0, 1, 20 | 2, 1, 2, 16 | .18 |
| **Ethnicity<sup>b</sup>** |  |  |  |  |  |
| *No. Hispanic/Latino* | 1 | 1 | 1 | 3 | .50 |
| **FSIQ<sup>c</sup>** |  |  |  |  |  |
| *range* | [90 - 147] | [88 - 124] | [67 - 131] | [63 - 115] |  |
| *mean* | 117.96 (16.46) | 106.5 (10.46) | 104.95 (14.76) | 95.43 (13.73) | .00002 |
| **Motion<sup>d</sup>** | 0.23 (0.11) | 0.28 (0.15) | 0.25 (0.11) | 0.32 (0.23) | .21 |
| **Handedness<sup>e</sup>** |  |  |  |  |  |
| *No. L,R* | 3, 19 | 2, 20 | 2, 20 | 2, 18 | .95 |

a: African American, Asian, Biracial, Caucasian, b: FSIQ: WISC-IV full-scale IQ, c: Mean framewise displacement, d: Number of children with left, ambidextrous, right, handedness, TD: typically developing, ASD+ADHD: ASD with comorbid ADHD

**Table S2.** ‘EF subgroup sample’ demographics

|  | EF subgroups |  |  | *P*<br>value |
| --- | --- | --- | --- | --- |
|  | Above Average<br>*n*=43 | Average<br>*n*=43 | Impaired<br>*n*=43 |  |
| **Sex** | 30 M/13 F | 35 M/8 F | 35 M/8 F | .33 |
| **Age** |  |  |  |  |
| *range* | [8.42 - 12.58] | [8.00 - 12.75] | [8.00 - 12.92] |  |
| *mean* | 10.19 (1.02) | 10.20 (1.37) | 10.04 (1.33) | .81 |
| **Race<sup>a</sup>** | 7, 2, 6, 28 | 3, 1, 6, 33 | 5, 1, 4, 33 | .79 |
| **Ethnicity, No. Hispanic/Latino** | 2 | 0 | 7 | .009 |
| **FSIQ<sup>b</sup>** |  |  |  |  |
| *range* | [90 - 142] | [84 - 147] | [63 - 129] |  |
| *mean* | 116.86 (12.44) | 111.43 (14.54) | 103.72 (13.78) | .00008 |
| **Motion<sup>c</sup>** | 0.24 (0.13) | 0.23 (0.12) | 0.30 (0.20) | .06 |
| **Handedness<sup>d</sup>, No. L,A,R** | 3, 1, 39 | 5, 0, 38 | 4, 0, 38 | .65 |
| **No. (%) in EF group** |  |  |  |  |
| TD | 41 (95%) | 15 (35%) | 0 |  |
| ADHD | 1 (2%) | 16 (37%) | 19 (44%) |  |
| ASD | 1 (2%) | 10 (23%) | 6 (14%) |  |
| ASD+ADHD | 0 | 2 (5%) | 18 (42%) |  |

a: African American, Asian, Biracial, Caucasian, b: FSIQ: WISC-IV full-scale IQ, c: Mean framewise displacement, d: Number of children with left, ambidextrous, right, handedness, ASD+ADHD: ASD with comorbid ADHD

**Table S3.** Medication status

| Table S3. Medication status |  |  |  |
| --- | --- | --- | --- |
|  | None | Non-stimulant | Stimulant |
| *EF subgroups* |  |  |  |
| Above average EF | 43 (100%) | 0 (0%) | 0 (0%) |
| Average EF | 24 (57%) | 5 (12%) | 13 (31%) |
| Impaired EF | 22 (52%) | 2 (5%) | 18 (43%) |
| *Diagnostic groups* |  |  |  |
| TD | 42 (100%) | 0 (0%) | 0 (0%) |
| ADHD | 14 (33%) | 0 (0%) | 29 (67%) |
| ASD | 24 (56%) | 7 (17%) | 11 (26%) |
| *Diagnostic comorbid groups* |  |  |  |
| TD | 22 (100%) | 0 (0%) | 0 (0%) |
| ADHD | 6 (29%) | 0 (0%) | 15 (71%) |
| ASD | 15 (68%) | 4 (18%) | 3 (14%) |
| ASD+ADHD | 11 (55%) | 2 (10%) | 7 (35%) |
| Distribution of participants who were taking no medication, taking non-stimulant psychotropic medication or taking stimulant medication at the time of study enrollment. |  |  |  |

**Figure S1.**
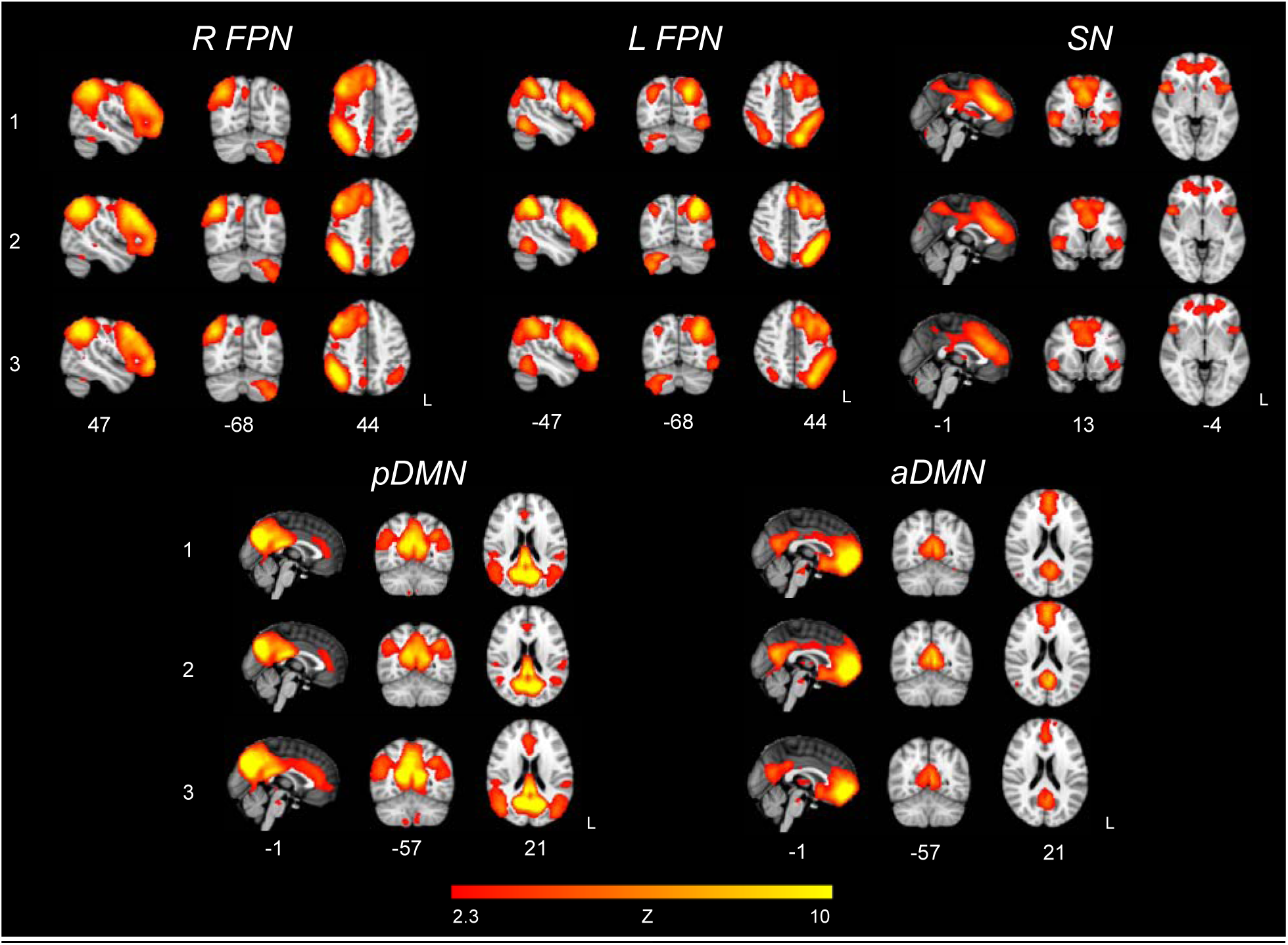
ICA components for 5 major cognitive networks across subsamples. The five major cognitive networks (R FPN, L FPN, SN, posterior DMN, anterior DMN) were estimated in three separate group ICAs for each subsample of interest: 1) EF subgroups, 2) Diagnostic groups, 3) Diagnostic comorbid groups. This figure qualitatively illustrates the similarities between estimated components for each subsample.

